# Data-Driven Extraction of a Nested Structure of Human Cognition

**DOI:** 10.1101/105403

**Authors:** Taylor Bolt, Jason S. Nomi, B. T. Thomas Yeo, Lucina Q. Uddin

## Abstract

Decades of cognitive neuroscience research have revealed two basic facts regarding task-driven brain activation patterns. First, *distinct* patterns of activation occur in response to different task demands. Second, a superordinate, dichotomous pattern of activation/de-activation, is commonly observed across a variety of task demands. We explore the possibility that a hierarchical model incorporates these two observed brain activation phenomena into a unifying framework. We apply a latent variable approach, exploratory bi-factor analysis, to a large set of brain activation patterns to determine the potential existence of a nested structure of factors that underlies a variety of commonly observed activation patterns. We find that a general factor, associated with a superordinate brain activation/de-activation pattern, explained the majority of the variance (52.37%). The bi-factor analysis also revealed several sub-factors that explained an additional 31.02% of variance in brain activation patterns, associated with different manifestations of the superordinate brain activation/de-activation pattern, each emphasizing different contexts in which the task demands occurred. Importantly, this nested factor structure provided better overall fit to the data compared with a non-nested factor structure model. These results point to domain-general psychological process, representing a ‘focused awareness’ process or ‘attentional episode’ that is variously manifested according to the sensory modality of the stimulus and degree of cognitive processing. This novel model provides the basis for constructing a biologically-informed, data-driven taxonomy of psychological processes.

---

A central goal of cognitive neuroscience is to understand cognitive processes through the analysis and examination of neuroimaging data. In the past two decades, researchers have attempted to delineate the brain activation patterns associated with various task demands to provide a neural grounding of cognitive processes. Thus far, hundreds of task fMRI studies have purported to demonstrate differential brain activation patterns associated with working-memory (Rypma and D’Esposito, 1999; Pessoa et al., 2002; Curtis and D’Esposito, 2003), attention (Brefczynski and DeYoe, 1999; Kanwisher and Wojciulik, 2000; Corbetta and Shulman, 2002), inhibitory control (Liddle et al., 2001; Menon et al., 2001), multi-sensory integration (Driver and Noesselt, 2008; Hong et al., 2009), affective regulation (Bechara et al., 1994; Damasio et al., 2000; Singer et al., 2004), and other cognitive processes. In contrast to this emphasis on linking individual cognitive processes with unique brain activation patterns, other researchers have demonstrated the existence of a dominant, canonical pattern of brain activation/de-activation that occurs across a variety of task demands and associated cognitive processes, sometimes referred to as ‘task-positive’ activation and ‘task-negative’ de-activation (Fox et al., 2005a; Toro et al., 2008; Duncan, 2010; Fedorenko et al., 2013; Hugdahl et al., 2015a). This canonical pattern of brain activation/de-activation involves an increase in activity in a collection of brain regions, often including the lateral prefrontal cortex, superior parietal cortex, and posterior medial prefrontal cortex, along with a corresponding decrease in midline regions, including the medial prefrontal cortex and the precuneus/posterior cingulate cortex. Researchers have argued that this canonical brain activation pattern represents a domain-general cognitive process (Duncan, 2010; Fedorenko et al., 2013; Hugdahl et al., 2015a) activated in response to the presence of external stimuli, irrespective of the content of the stimuli.

These observations give rise to two tenets. First, that the brain produces distinct patterns of activation in response to different task demands. Second, that the brain produces a singular, superordinate pattern of activation/de-activation across a variety of task demands. Recent work has aimed to reconcile this apparent dichotomy by exploring the extent to which different tasks share common and distinct psychological processes (Poldrack et al., 2009; Barrett and Satpute, 2013; Krienen et al., 2014). We hypothesized that these two streams of research, rather than contradictory, point to the existence of a *nested* structure of brain activation patterns underlying human cognition. In particular, we hypothesized that an overarching domain-general cognitive process, represented by the canonical brain activation/de-activation pattern, is present across all task demands, in agreement with previous literature, but presents in different manifestations depending on cognitive processes unique to the type of task demands. Thus, cognitive processes unique to different types of task demands are distinguished by differential sub-types of the canonical task-positive/task-negative pattern.

We tested this hypothesis with a novel application of a bi-factor analytic model applied to unthresholded task-activation maps from a large of sample of published fMRI studies (**Figure 1**). The use of unthresholded brain activation maps provides an advantage over previous studies (Toro et al., 2008; Smith et al., 2009; Lenartowicz et al., 2010; Bertolero et al., 2015; Yeo et al., 2015) examining task-general brain activation patterns that have used activation-coordinate databases, such as the BrainMap database (Fox and Lancaster, 2002) or Neurosynth (Yarkoni et al., 2011). Coordinate-based analyses are inherently limited due to their reduction of full statistic images to peak-activation coordinates (Salimi-Khorshidi et al., 2009; Poldrack and Yarkoni, 2016), and at present, there are no universal standards for reporting activation coordinates (Wager et al., 2007). The use of ‘full-information’ activation images, rather than peak-activation coordinates, allows for more accurate assessments of the covariance between any two activation maps, and a wider variety of analytic approaches to explore the associations among a group of activation maps, such as the bi-factor analysis approach applied in this study. Utilizing a largescale dataset of unthresholded brain activation maps from several task domains provided through the Human Connectome Project (Barch et al., 2013) and NeuroVault database (Gorgolewski et al., 2015), this study directly tested the hypothesized nested or bi-factor organization of brain activation patterns. In addition, we directly compared the overall fit to the collection of brain activation maps of a nested factor model of brain activation patterns to a non-nested factor model of brain activation pattern.

**Figure 1.**
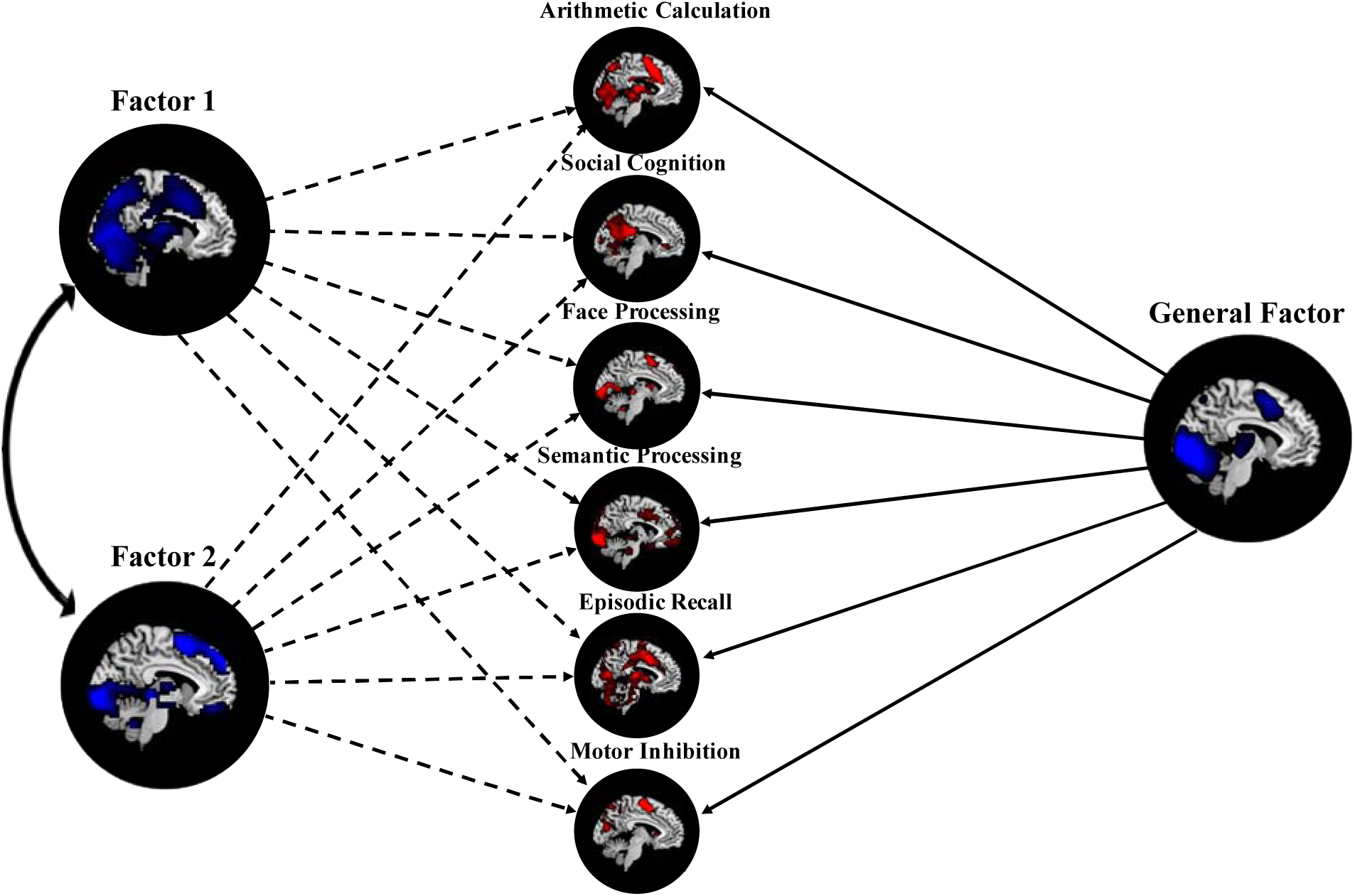
Illustration of Exploratory Bi-Factor Analysis Approach. In this particular application of the bi-factor analytic model, each task-activation map, representing the activation (in red) associated with a particular task state (e.g. social cognition or semantic processing), are modeled as arising from direct effects of a *single* latent general factor and various latent sub-factors (e.g. Sub-Factor 1 and Sub-Factor 2). In addition, sub-factors may also be correlated with one another, indicated by the double-headed arrow. These latent factors can also be represented in brain space (represented in blue) through the computation of scores associated with factors.

## Materials and Methods

### HCP and NeuroVault Inclusion and Exclusion Criteria

Neuroimaging data from 208 unrelated, healthy, right-handed adults (Mean age = 28.61 years (SD: 3.85, range: 22 – 36); 103 female) made available through the Human Connectome Project (HCP) 2014 release were used for this study. These were the maximum number of unrelated subjects in the 2014 release (Barch et al. 2013). For each subject, approximately 7 task scans (note not all subjects had all task scans) were used for analysis. Here we refer to the tasks as the working-memory task, gambling task, motor task, language task, social task, relational task, and emotion task.

Eighty-six task contrasts for each subject were made available through the Human Connectome Project (HCP) 2014 release. Only those task contrasts that were independent of other contrasts (i.e. non-redundant) and explicitly modeled BOLD activation in response to block or event-related task onsets were included in the study (referred to as ‘brain activation maps’), leaving a total of 24 contrasts. In particular, many contrasts were negative contrasts that model de-activations associated with block onsets. These types of contrasts are perfectly correlated/anti-correlated (negative contrasts) with the contrast(s) of interest, and would prevent the estimation of the factor solution. Subtraction contrasts, or contrasts computed by a subtraction between two brain activation contrasts, were not included in the study because the study was explicitly interested in identifying factors underlying task-activation/de-activation patterns, rather than *relative* activation/de-activation between modeled contrasts. Thus, these types of task contrasts were not included in the analysis. Because the HCP provided *participant-level* brain activation maps, a standard group-level estimation procedure was used to estimate *group-level* brain activation maps (described below) from the participant-level maps.

At the time of the analysis (August 2016), the NeuroVault database (Gorgolewski et al., 2015) contained 369 publicly shared collections of images, where each collection contained all of the images from a single study. A collection in the NeuroVault database may contain unthresholded fMRI and PET statistic images, functional and structural parcellations, and anatomical atlases. Unthresholded fMRI group-level task contrasts were used for this study. In particular, group-level task contrast images representing average BOLD activity in response to blocked or event-related task onsets and offsets were of interest (referred to as ‘brain activation maps’). Excluded images were images derived from subtraction contrasts, meta-analyses, functional connectivity analyses (seed-based, Independent Components Analysis, etc.), multivariate pattern analyses (MVPA), behavioral correlation analyses, group comparison analyses, and any analyses of patient or clinical populations. To ensure the brain activation maps meeting this criteria were of sufficient quality, only studies using 20 or more participants were included (Desmond and Glover, 2002). Eighty-seven brain activation maps from 17 NeuroVault collections met these criteria and were included for preprocessing. Included in this total are two brain activation maps from the UK Biobank study (Miller et al., 2016) (further references to ‘NeuroVault’ brain activation maps include these two activation maps).

### Processing of HCP and Neurovault Data

For the HCP task data, the participant-level volume-based analyses were used to estimate group-level brain activation maps. The preprocessing of this task data involved gradient distortion correction, motion correction, registration to the Montreal Neurological Institute (MNI) template (MNI152 space), and grand-mean intensity normalization. In addition, spatial smoothing was applied to each task scan using an unconstrained 3D Gaussian kernel of FWHM=4mm. The details of the minimal preprocessing pipeline are described in Glasser et al. (2013). Session-level analyses were carried out within each task for both encoding directions and are described by Barch et al. (2013): session-level activity estimates were computed using the general linear model (GLM) implemented in FSL’s FILM (FMRIB’s Improved Linear Model) with autocorrelation correction (Woolrich et al., 2001). Predictors for each task contrast were convolved with a double gamma canonical hemodynamic response function (Glover, 1999). To account for slice-timing differences and variability in the HRF delay across regions, temporal derivatives of each predictor were added as a confound term to the GLM. Timeseries were then highpass filtered with a cutoff of 200 seconds and prewhitened within FILM to correct for autocorrelations in the timeseries. The two session-level activity estimates for each encoding direction were then combined using a fixed-effects GLM analysis using FSL’s FEAT to estimate average effect participant-level estimates. Group-level mixed-effect analyses treating subjects (n=208) as a random effect were conducted for each sample with the participant-level fixed-effect estimates using FSL’s FLAME (FMRIB’s Local Analysis of Mixed Effects) to estimate the average effects of interest across the sample (Woolrich et al., 2004). The resulting unthresholded group brain activation maps (z-statistic images) were used for further analyses.

Most of the brain activation maps in the sample chosen from NeuroVault were *t-statistic* images and were thus, converted to *z-statistic* images for normalization across activation maps. In order to ensure the group brain activation maps from the NeuroVault database were adequately comparable, several steps were taken to reduce inter-scanner and inter-subject variability. First, all NeuroVault maps were re-registered to a 2mm MNI template using FSL’s FLIRT (Jenkinson and Smith, 2001). Three of the 87 brain activation maps from the NeuroVault database failed to align to the MNI template using this process, and were excluded from further analyses, leaving 108 (24 HCP and 84 NeuroVault maps) brain activation maps overall for the analysis. To further reduce inter-study variability and ensure comparability across both NeuroVault and HCP maps, all brain activation maps were resampled to 3mm voxel size (voxel resolution of all maps ranged from 2-4mm) and were spatially smoothed with a 6mm FWHM Gaussian kernel.

### Conjunction Analysis

For a preliminary analysis of the total set of 108 brain activation maps (*z*-statistic images), a simple conjunction analysis was performed. All brain activation maps were thresholded at *z*=2.3 and *z*= −2.3 (*p* <.01) to examine the conjunction of activation and deactivation across the maps, respectively, and then binarized, such that suprathreshold activation/de-activation areas were replaced with a value of one. All activation and de-activation images were then summed across all maps to form activation and de-activation conjunction images, respectively.

### Exploratory Bi-Factor Analysis

The goal of all common factor analytic models, including the bi-factor model, is to explain the common variability among a group of observed variables (i.e. brain activation maps) in terms of a smaller set of unobserved or latent variables, called factors. The brain activation maps are modelled as causally influenced by, or indicators of, their respective factors(MacCallum, 2009). Thus, an implicit model of the relationship between the observed variables and unobserved factors in factor analysis is assumed (i.e. model-based analysis) in factor analysis, distinguishing this approach from principal components analysis (PCA), a similar approach. Specifically, the common factor model, including the bi-factor model, represents each observed variable as a linear combination of *common* and *unique* factors:

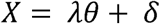

Where *X* is the *n (observations) × p (variables)* vector of scores on the observed variables; λ is the *p (variables) × m (common factor)* estimated matrix of factor loadings describing the association between each variable and each common factor; θ is the scores on the common factors; and δ is the *m* (unique factors) × 1 matrix of unique factor scores (“measurement error”) or the variance unexplained by the common factors. In the bi-factor model, restrictions (described below) are placed on the factor loading matrix (λ) so as to model a general factor in one column of the matrix and sub-factors in the remaining columns. In the application of the model to the current data, each brain activation map corresponding to the average BOLD activation/de-activation in response to particular processes (e.g. arithmetic calculation) or stimuli (e.g. angry faces) is modeled as a linear combination of a general latent factor that causally influences all variables and nested sub-factors that causally influence subsets or groupings of the brain activation maps.

One major difference between the latent variable approach used in this study and other common data-driven fMRI approaches, such as the standard fMRI independent components analysis approach (Calhoun et al., 2008; Smith et al., 2009), is that the factor analytic model in this study is estimating relationships *among* brain activation maps rather than estimating relationships among voxels within the brain activation maps. The primary goal of this study was to estimate underlying factors that explain the common variance among the brain activation maps, rather than the common variance among voxel activity patterns. Thus, the latent variables in this approach are computed as linear combinations of brain activation maps rather than voxels. This is made possible by the use of unthresholded statistical images that allow for more accurate assessments of covariance between any two maps. In addition, we were interested in *representative* patterns of task activation/de-activation that explain as much variance as possible among the brain activation maps, thus, the factor analytic model extracts underlying factors that best explains the covariance in the data, rather than estimating possible independent sources underlying the associations among the brain activation maps (i.e. independent components analysis).

The exploratory bi-factor analysis approach in this study proceeded in three stages: 1) a data reduction stage, 2) an initial extraction phase, and 3) a bi-factor analysis phase. The extraction and rotation phases were conducted using the exploratory factor analysis program in Mplus software (Muthén and Muthén, 2011).

### Data Reduction Stage

One assumption of the factor analysis models is independence of observations. Because of the strong spatial dependencies among voxels (n=80,090) within a brain activation map, this assumption is unlikely to hold. Thus, a high-resolution, data-driven parcellation (n=950; **Figure 2**) of the cortex was used (Craddock et al., 2012), and voxel z-scores within the ROI’s of the parcellation were averaged together. This high-resolution parcellation included only gray matter cortical and sub-cortical regions of the brain, excluding brainstem regions. The brainstem was excluded in this study because it is often difficult to obtain reliable signal from this area (Brooks et al., 2013). In order to ensure the parcellation adequately represented the data, we conducted a between-ROI analysis of variance (ANOVA) for each brain activation map to determine how much of the variance in each map was accounted for by between-ROI variance as opposed to within-ROI (error) variance. The results of the between-ROI ANOVA revealed that the ROI parcellation accounted for a substantial majority of variance in the data across all brain activation maps (Mean ROI_B/W_ = 0.829%, range: 0.665 – 0.882%).

**Figure 2.**
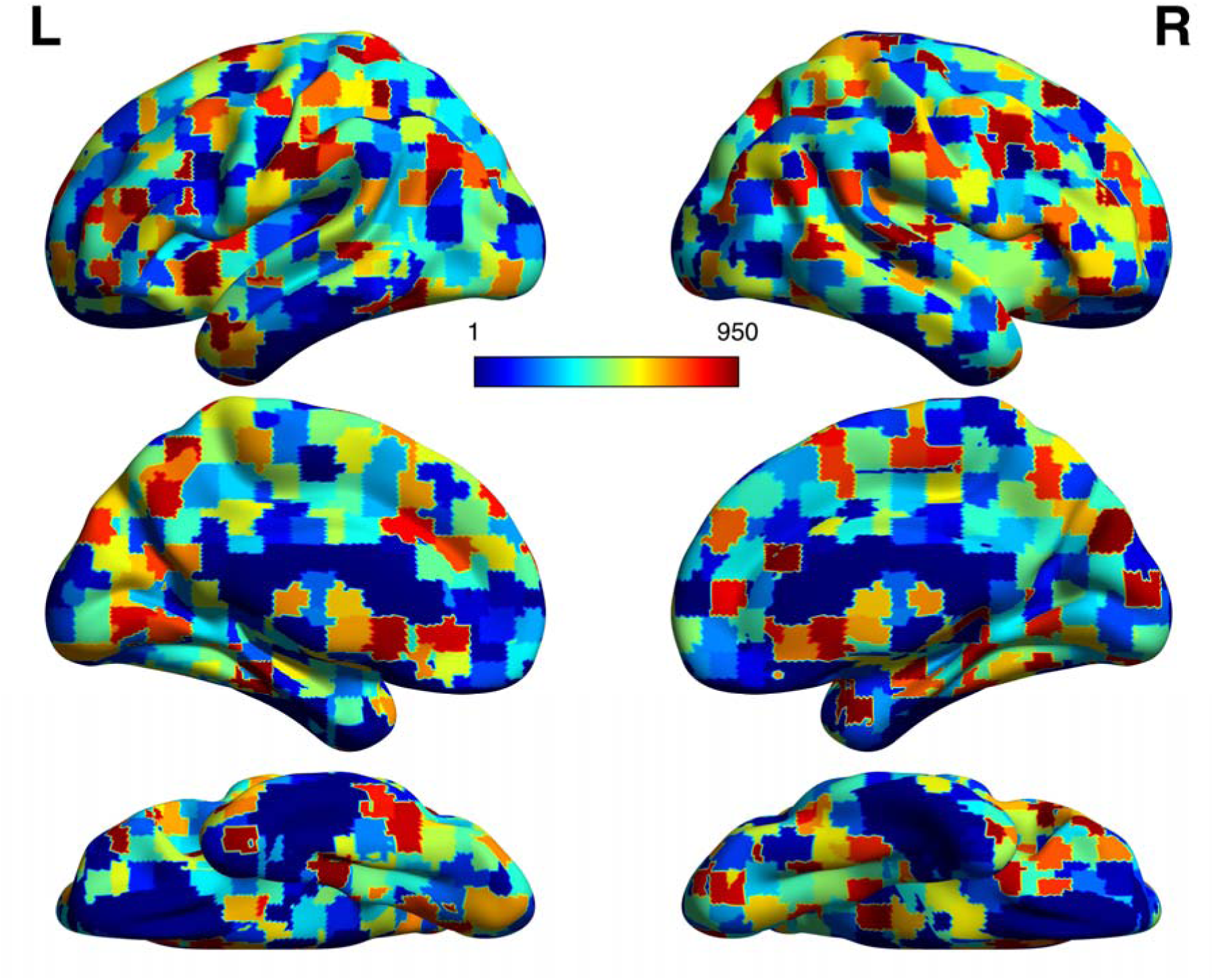
Craddock High-Resolution Parcellation (n=950). The a priori parcellation used as an initial data reduction before the bi-factor analysis. The parcellation was computed using a spatially constrained spectral clustering technique to define cortical ROIs with homogenous functional connectivity relationships (Craddock et al., 2012). ROIs are differentiated by integer values (1–950) and visualized on a surface MNI template.

### Initial Extraction Phase

The ROI values for each brain activation map were then vectorized and placed into a matrix, where ROIs represent (n=950) observations (i.e. rows) and columns represent variables (n=108). Thus, the data matrix *X* described above corresponds to the *n* ROI values (950) × *p* activation map (108) matrix. In the initial extraction phase of the factor analysis approach, the number of factors is estimated. To estimate the number of factors in the data we used the parallel analysis method (O’connor, 2000). This method involves extracting components and their associated eigenvalues (analogous to the amount of explained variance) through principal component analysis (PCA) applied to permutations (n=10,000) of the original data matrix (*X*) using column-wise reshuffling of the original data matrix. The mean permuted eigenvalues are then compared to the component eigenvalues extracted from the original data matrix (*X*) to determine the number of factors in the solution. In particular, the estimated number of factors were the number of components with eigenvalues greater than the mean permuted component eigenvalues.

### Bi-Factor Analysis Phase

After the number of factors was estimated, a maximum likelihood estimator with robust standard errors (MLR) was used to estimate the bi-factor model. Maximum likelihood estimation operates through an iterative search over parameters (factor loadings matrix and specific variances) that maximize a log-likelihood function, which measures the likelihood of the following parameters producing the sample variance-covariance matrix. However, the original maximum likelihood estimator assumes a multivariate normal distribution of the variables in estimation of model parameters, and a multivariate test of normality (Royston, 1983) (*p* < 0.0001) indicated that the joint distribution of the data did not exhibit normality. Thus, a robust maximum likelihood estimation procedure was chosen to correct for standard errors to correct for departures from non-normality. For detailed information on the implementations of these algorithms in Mplus, see (Muthén and Muthén, 2011). In order to examine possible bi-factor or nested structure among the brain activation maps, a bi-factor rotation criteria was applied to the factor loading matrix (λ) of the initial solution (Jennrich and Bentler, 2012). Importantly, the use of rotation procedures does not alter the fit of the model to the data (MacCallum, 2009). The bi-factor rotation criteria attempts to rotate the initial factor loading matrix towards a loading matrix with strong loadings of the first factor (general factor) and perfect cluster structure for the loadings on the remaining factors (sub-factors). There are two types of bi-factor rotation criteria, orthogonal and oblique rotation criteria (Jennrich and Bentler, 2012). An orthogonal rotation constrains the factors to be uncorrelated, while an oblique rotation allows the factors to correlate with each other. Because orthogonal rotations constrain the factors to be uncorrelated and are generally not recommended without an *a priori* hypothesis of independence for the factors (Costello & Osborne, 2005; MacCallum, 2009), we used the bi-factor *geomin* rotation criteria, a commonly used oblique rotation criteria (Hendrickson and White, 1964). The bi-factor *geomin* rotation criteria maximizes loadings for all maps on the first column (corresponding to the general factor) of the factor loading matrix, and minimizes a measure called *complexity* (the number of nonzero elements in the corresponding row of the factor loading matrix) for all activation maps on the remaining columns of the factor loading matrix, which has the effect of enforcing sparsity on the loadings in the matrix. The output of the rotation procedure is a new ‘rotated’ factor loading matrix (i.e. pattern matrix), representing the correlation between each brain activation map and each factor, controlling for its association with the other factors. The pattern of factor loadings on each factor was then used to interpret the types of task demands/cognitive processes associated with each factor. Significance estimates are also estimated for each activation map-factor loading pair from the estimated robust standard errors (derived from the MLR estimation procedure). The significance estimates for the factor loadings were not corrected for multiple comparisons (*p<.05*), given the results implied by the factor analysis (10 factors underlying the substantial majority of the variance in the data) imply strong dependencies among the brain activation maps.

The MLR estimation procedure also produces explained variance estimates for each factor, and *communality* estimates for each brain activation map. The explained variance estimates represent the variance among the activation maps that is explained by each factor. However, these estimates for each factor are approximate in an oblique solution where factors are correlated, and are more accurate to the degree that factor is orthogonal (statistically uncorrelated) with the other factors in the solution. Thus, all estimates of explained variance associated with each factor are approximate, excluding the general factor, which is orthogonal to the sub-factors. The *communality* estimate is the proportion of variance in that variable accounted for by the extracted factors, and represents the degree to which that brain activation map is represented in the solution.

### Computation of Factor Scores

In order to compute a brain region’s (i.e. each ROI of the high-resolution parcellation) standardized ‘score’ for each factor, we used a least-squares estimation method, known as the Bartlett method (Bartlett, 1937; Lawley and Maxwell, 1962). The Bartlett method of factor score computation is similar to the ordinary least squares approach, where the factor score (θ vector in *Equation 1*) at each region is estimated from the observed *X* data matrix and λ matrix of factor loadings, but the added condition that the *communality* estimates (described above) weight each activation map from the factor solution. Thus, brain region values from activation maps that are better captured by the factor solution contribute more to the factor score at that brain region (Bartlett, 1937). The resulting factor scores for each region were then mapped back to the brain, smoothed using a 6mm FWHM Gaussian kernel for smoother boundaries among neighboring regions of the brain, and visualized on an MNI152 surface template using BrainNet Viewer software (Xia et al., 2013). The resulting brain maps correspond to each factor’s representation in standard brain space.

### Comparison of Alternative Factor Structures with Bi-Factor Model

In order to test whether alternative factor models fit the data equally well, we used a confirmatory factor analysis (CFA) approach that measured the overall fit of two alternative factor models compared to the bi-factor model. CFA operates in the opposite ‘direction’ from the exploratory factor analysis model used above, and involves the fitting of an *a priori* factor model, with variable-factor loadings specified by the user, to the data. From a path diagram perspective, this can be conceptualized as specifying *a priori* links from the factors to the brain activation maps. The overall fit of the *a priori* factor models to the data is assessed using a variety of global fit indices (Fan et al., 1999; Bentler, 2007; Iacobucci, 2010) that all measure the discrepancy between the covariance matrix implied by the factor model and the sample covariance matrix (covariance between the observed variables). Four global fit indices were used to compared the overall fit of each model to the data: root mean square error of approximation (RMSEA), comparative fit index (CFI), Tucker-Lewis index (TLI), and the standardized root mean square residual (SRMR). Lower values for the RMSEA and SRMR represent greater over all fit, and higher values for the CFI and TLI represent greater overall fit. As with the exploratory bi-factor analysis, both CFA models were analyzed in Mplus.

### Study/Sample-Specific Non-Nested Alternative Factor Model

One possible concern with combining together unthresholded statistic images across studies is that the majority of differences among the statistical images may be driven by sampling and scanner-site differences (Friedman et al., 2006). Overall, there were 19 collections of brain activation maps, but four collections consisted of one map and a factor with a single indicator, which was not identifiable (i.e. parameters cannot be uniquely estimated) (Brown, 2015), and were not included in the CFA for the standard non-nested alternative exploratory factor analysis model and bi-factor model described below. Thus, the non-nested factor model included 15 orthogonal factors (no covariance between factors), corresponding to 15 collections (13 Neurovault collections, HCP, and BioBank) of brain activation maps. Three factors (corresponding to three collections) consisted of two activation maps and were not identifiable (because of the independence of the factor), and were thus allowed to arbitrarily co-vary with the HCP factor, the largest factor, for identification. Note, the adding of parameters to be estimated, however arbitrary, does not decrease, but improves overall model fit.

### Standard Exploratory Factor Analysis Non-Nested Alternative Factor Model

In order to compare the data-driven nested bi-factor model to an alternative data-driven non-nested factor model, we performed a standard non-nested exploratory factor analysis with a *promax* rotation (Hendrickson and White, 1964; Cureton and Mulaik, 1975) using the same MLR estimation procedure used for the bi-factor analysis. Ten non-nested factors were chosen to be estimated, the same as the number of factors in the bi-factor model. The resulting rotated factor loadings were then used to construct a non-nested factor model. Specifically, each activation map was specified as loading on the factor to which it had the strongest loading. Eight non-orthogonal (allowed to co-vary) factors were specified in the factor model, as one factor did not have a single strongest activation map loading and the other factor had only one strongest activation map loading, and was not identifiable; to keep the one activation map, (‘Simon_Incongruent_Incorrect’) in the factor model, it was specified as loading on the factor to which its paired activation map (‘Simon_Congruent_Incorrect’) had the strongest loading.

### Nested Bi-Factor Model

The results from the exploratory bi-factor analysis were used to construct a confirmatory bi-factor model. As described in the results, the bi-factor model included ten factors, a general factor, and nine sub-factors. For each brain activation map, two factor loadings were specified: a factor loading on the single, general factor, along with a single sub-factor, corresponding to its strongest sub-factor loading from the exploratory bi-factor analysis.

## Results

### Conjunction Analysis

To test whether the full set of 108 brain activation maps exhibited the commonly observed superordinate, task-positive and task-negative activation pattern, we performed a conjunction analysis by summing binarized activation and de-activation estimates across all maps. The results (**Figure 3**) for both activation and de-activation maps correspond closely to the classic task-positive and task-negative activity described in previous neuroimaging studies(Raichle et al., 2001; Dosenbach et al., 2007; Duncan, 2010; Fedorenko et al., 2013). The strongest areas of overlap in the brain-activation conjunction analysis included the posterior medial prefrontal cortex (pMPFC), anterior insula (AI), superior parietal cortex (SPC), dorsolateral prefrontal cortex (DLPFC), and visual network areas. Activation overlap was strongest in the pMPC and in the fusiform area of the visual network, with some voxels within these areas active across 90% (97 out of 108) of all activation maps. The highest areas of overlap in the task-deactivation conjunction were in the default mode network, comprised of the medial prefrontal cortex (MPFC), the precuneus/posterior cingulate cortex (precuneus/PCC) and the bilateral inferior parietal cortices (IPC). Interestingly, average overlap was higher in the task-activation conjunction (*Max*_*Overlap*_=97) compared with the task-deactivation conjunction (*Max*_*Overlap*_=81), indicating that the pattern of task-related de-activation was less consistent (i.e. more variable) compared with the pattern of task-related activation.

**Figure 3.**
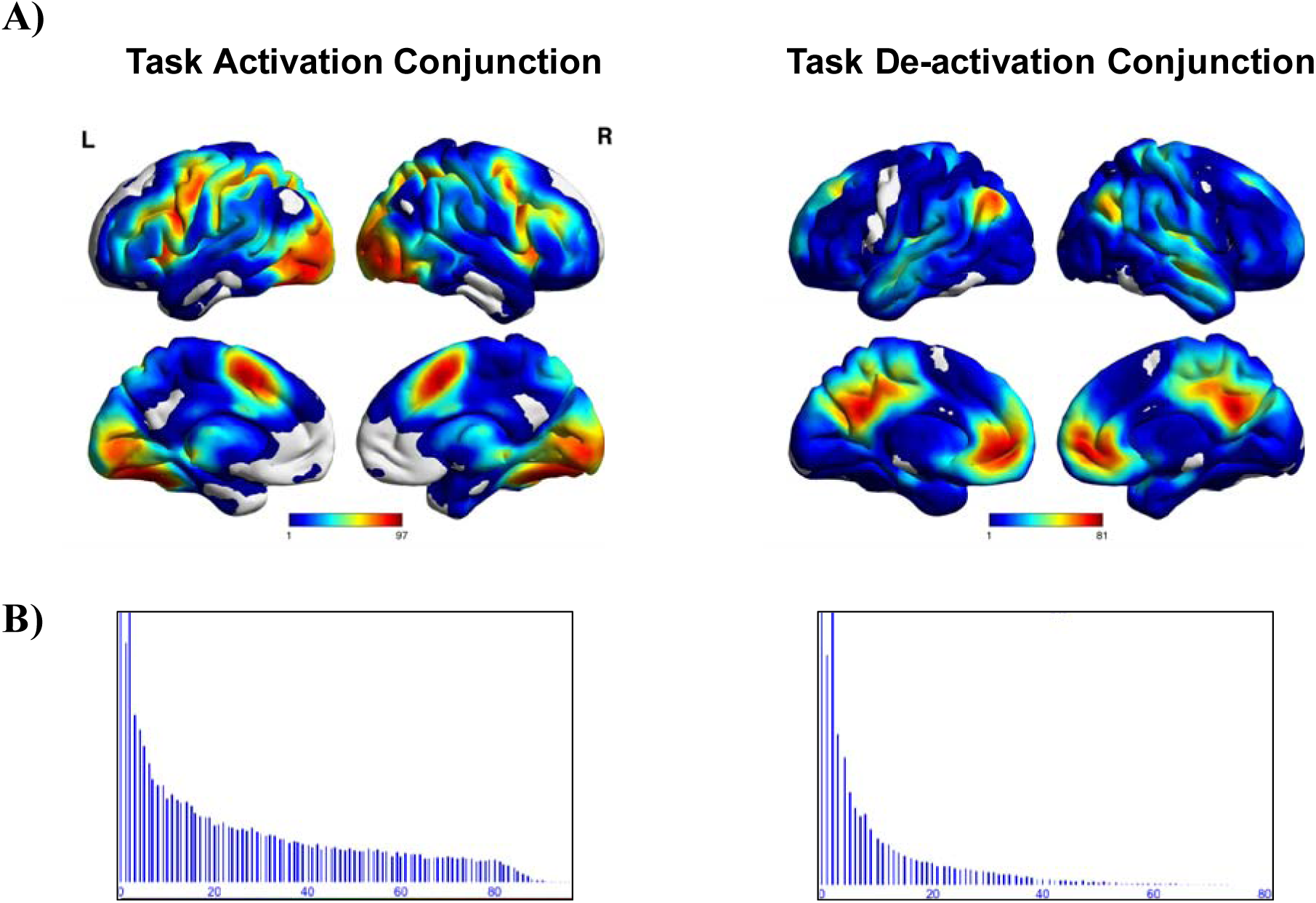
Conjunction Analysis of All Activation Maps. The results of the conjunction analysis examining the consistency/overlap of the task-activation/de-activation across all activation maps. **A)** Conjunction maps visualized on an MNI surface template. Values for each voxel represent the amount of times that voxel was activated or de-activation across all 108 activation maps. **B)** Histogram displaying the frequency distribution of the consistency/overlap values for all voxels. The distribution follows an exponentially decreasing curve with the majority of voxels observed to have low consistency values. As can be seen from the conjunction maps and the histograms, greater consistency values (both average and max consistency values) are observed in task-activation, as opposed to task-deactivation patterns.

### Initial Extraction

In order to estimate the number of factors among the brain activation maps, a parallel analysis was conducted (**Figure 4**) that involved comparison of mean ‘random’ component eigenvalues (analogous to the amount of explained variance) from permuted versions of the original data matrix to the component eigenvalues extracted from the original data matrix. The parallel analysis indicated that a 10 factor solution best represented the data, explaining a total of 83.39% percent of the variance among the activation maps.

**Figure 4.**
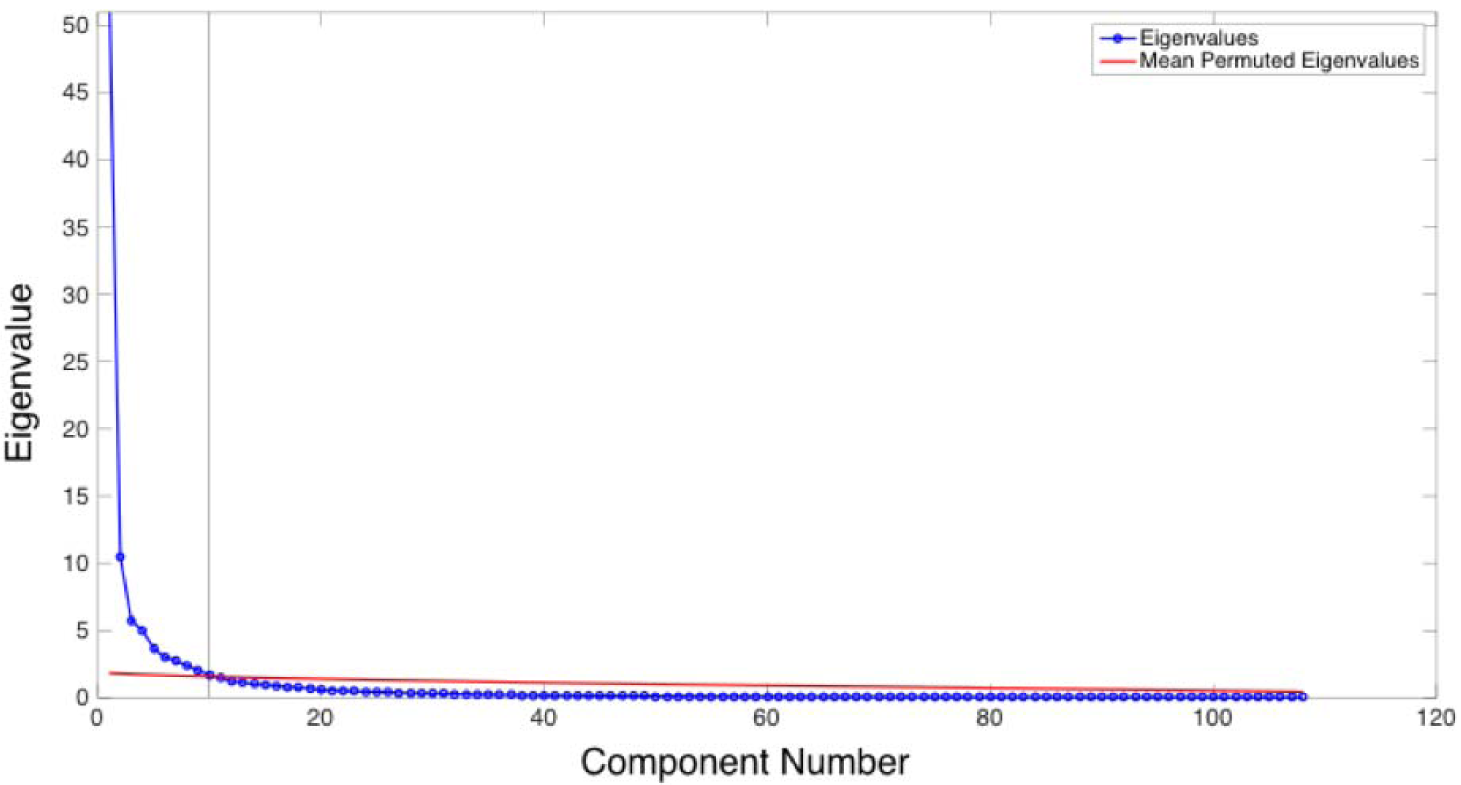
Parallel Analysis. A plot of the mean permuted eigenvalues (red) and the component eigenvalues of the original data matrix. The eigenvalue is plotted along the vertical axis and the component number is plotted along the horizontal axis. Component eigenvalues were greater than the mean permuted eigenvalues up to component 10 (indicated by the horizontal black line), indicating a ten factor solution was appropriate.

### Bi-Factor Analysis

A bi-factor model was then estimated from the data with a ten factor solution involving one single general factor and nine sub-factors. The communality estimates, representing the proportion of variance in each variable accounted for by the bi-factor solution (R^2^), indicated that the substantial majority of activation maps were captured fairly accurately in the bi-factor solution (R^2^ > 0.4) (Costello and Osborne, 2005). Remarkably, only four of the 108 activation maps were not adequately captured by the 10 factor solution, (R^2^ < 0.4). Examination of the activation/de-activation patterns of the low communality activation maps revealed that these maps deviated significantly from the dominant task-positive/task-negative pattern with activation localized to the motor cortex (activation maps associated with simple motor movements) (**Figure 5**). The pattern of factor loadings, representing the association between each activation map and each factor, was then examined for potential bi-factor or nested structure.

**Figure 5.**
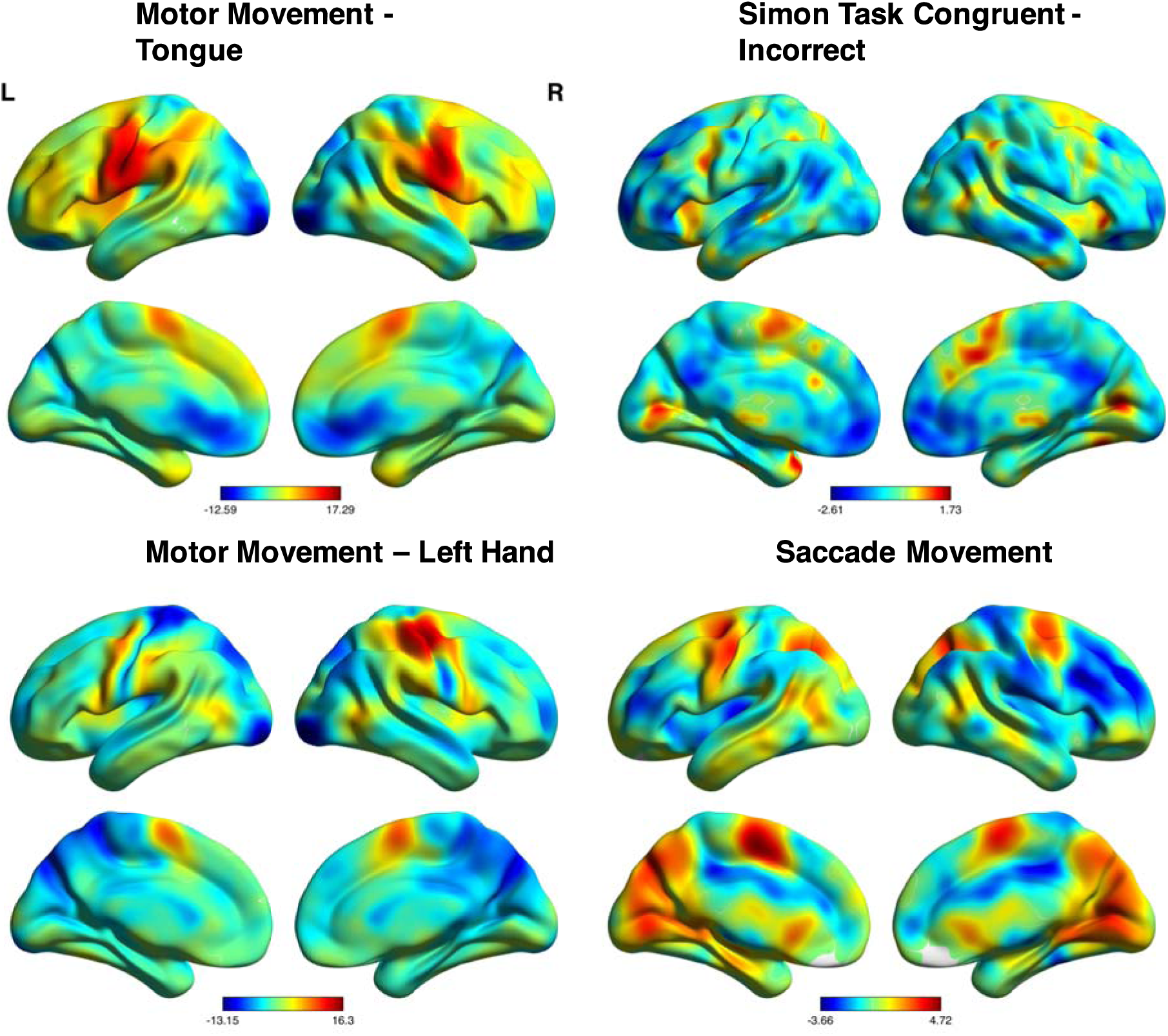
Brain Activation Maps with Low Communalities in Bi-Factor Solution. Z-statistic images for motor tongue movements, incorrect trials for the incongruent trial types of the Simon Task, motor left hand movements, and average saccade movement from a saccade conflict task. Examination of activity patterns from all four activation maps reveals that they deviate from the dominant pattern of activity in **Figure 2**.

### General Factor

As hypothesized, a general factor was prevalent in the data (**Figure 6**). In particular, the general factor accounted for the majority of the common variance among the brain activation maps (52.37%), and 94 out of 108 maps had statistically significant loadings on the general factor. The 14 brain activation maps that did not have significant loadings on the general factor were associated with simple motor movements (e.g. finger movements in response to visual cues) and processing of simple auditory stimuli (e.g. button presses in response to an auditory cue). The strongest factor loadings on the general factor included a variety of types of task demands and stimuli types: affective facial processing, visual working-memory, and visual pattern comparison. Factor score patterns (see *Methods and Materials*) in the cortex, representing the degree that each factor is present (positively or negatively) in each region of the brain, corresponded to the canonical task-positive/task-negative pattern observed in previous studies: positive scores (activation) in posterior medial prefrontal cortex (pMPFC), dorsolateral prefrontal cortex (DLPFC), and the superior parietal cortex (SPC), along with negative scores (de-activation) in the medial prefrontal cortex (MPFC), precuneus/posterior cingulate cortex (P/PCC), and inferior parietal cortex (IPC). Activation also extended into primary visual cortex (V1), higher-order visual object processing areas in occipito-temporal areas of the brain, such as the fusiform gyrus, as well as the primary motor cortex, possibly corresponding to the common experimental set-up of button responses from fingers of the right or left hand in response to stimuli.

**Figure 6.**
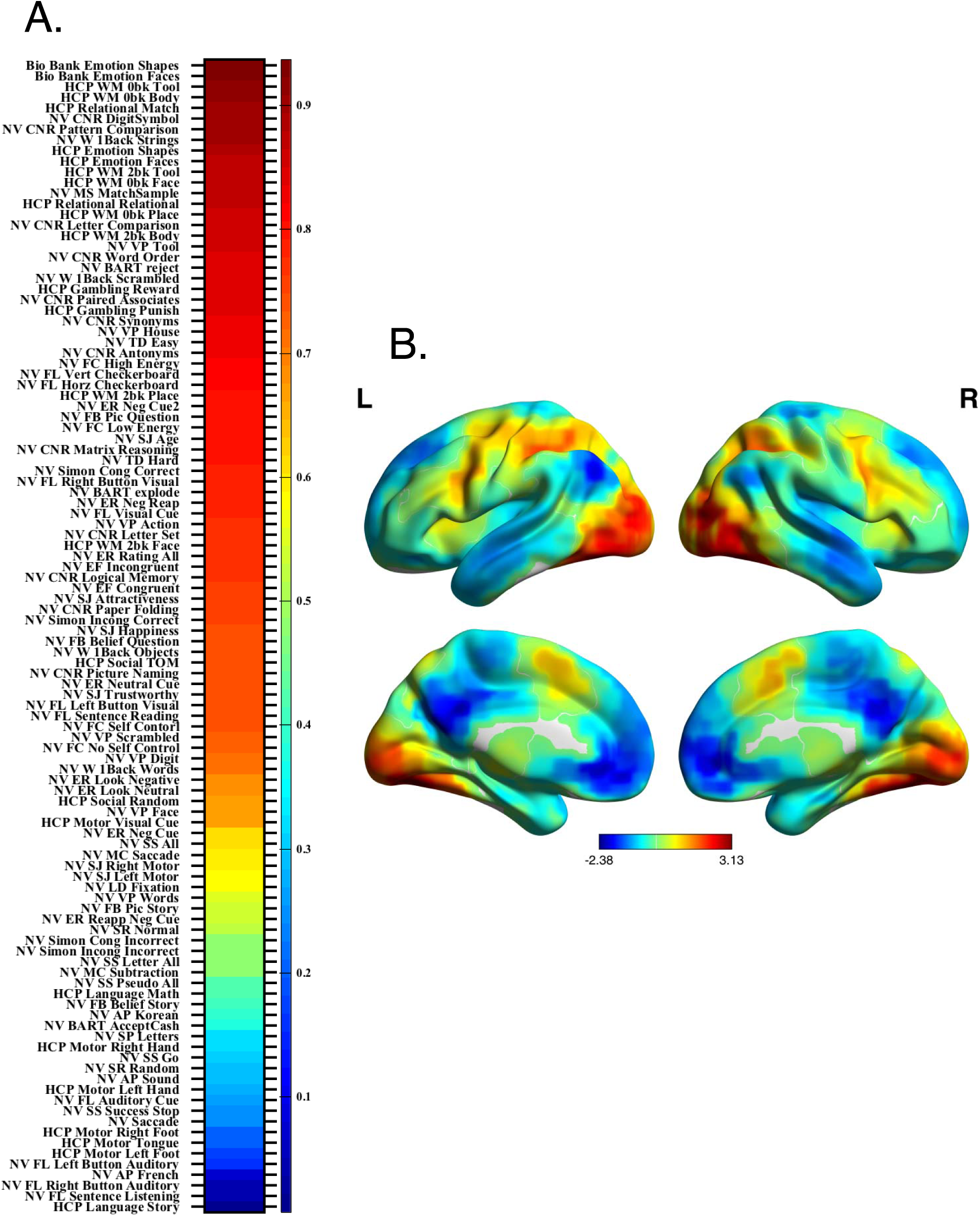
General Factor Loadings and Factor Score Visualization. (NV = NeuroVault; HCP = Human Connectome Project; CNR = Cognitive Neuroscience Lifespan Repository; AP Auditory Processing; BART = Balloon Analog Risk Taking; EF = Erikson Flanker; Episodic = Episodic Recall; ER = Emotional Regulation; FB = False Belief; FC = Food Choice; F-L = Functional Localizer; LD = Lexical Decision; LR = Logical Reasoning Task; MC = Mental Calculation; MS = Match to Sample; PC = Pattern Comparison; SD = Speech Detection; Simon = Simon Task; SM = Semantic Memory; SJ = Social Judgment; SR = Sentence Reading; SS = Stop Signal; TD = Temporal Discounting; TOM = Theory of Mind; VP = Visual Processing; W_1back = Word 1Back; WM = Working-Memory). **A)** The loading of each activation map on the general factor, which represents the association between each map and each factor. The activation map labels are displayed along the vertical axis and their associated loading values represented by a heat map are displayed to the right of the label (warmer colors represent stronger loadings and cooler colors represent weaker loadings). Out of 108 activation maps, 94 had significant loadings on the general factor. In particular, all activation maps with loadings stronger than the ‘HCP Motor Right Hand’ activation map (λ > 0.32) were found to be significant (*p* < 0.05), along with the ‘NV Saccade’ (λ = 0.24) and ‘NV SR Random’ (λ = 0.3) activation map. **B)** Factor scores for each region were placed in MNI coordinates and visualized on a surface MNI template (warmer colors represent positive scores and cool colors represent negative factor scores). The pattern of positive and negative scores for the general factor followed the canonical task-positive/task-negative activation/de-activation pattern.

### Sub-Factors

Factor loadings exhibited strong clustering across the nine distinct sub-factors (**Figure 7**), with at least four brain activation maps with strong loadings solely on each sub-factor. All sub-factors explained a similar small amount of variance in the data (~3%), with the exception of factor three that explained approximately 9% of the data. In addition, the sub-factors were relatively orthogonal (|r|=0.08; Range: −0.216 – 0.272), although the bi-factor model allowed for correlated sub-factors. Examination of the factor scores associated with each sub-factor revealed a variety of task-positive/task-negative patterns. Notably, positive scores in the pMPFC, encompassing the pre-supplementary motor area/supplementary motor area (pre-SMA/SMA) and dorsal anterior cingulate cortex (dACC), were observed across all sub-factors. High positive scores in the DLPFC and AI were also observed across many of the nine sub-factors as well, but with less consistency than the pMPFC. In agreement with the results of the conjunction analysis, the pattern of negative scores across the nine sub-factors was much less consistent. However, strong negative scores within at least one of the areas of the DMN (MPFC, P/PCC, and IPC) was observed across all sub-factors, with the exception of factor eight.

For initial categorization, sub-factors were grouped based on their pattern of factor scores into predominantly auditory-temporal, visual-parietal, and pre-frontal sub-factors. The auditory sub-factors included sub-factors three and four, and both exhibited strong positive scores in the primary auditory cortex (superior temporal gyrus). Outside of the primary auditory cortex, positive scores for sub-factor three were predominantly observed in regions of the pMPFC, AI, frontal poles and the sensorimotor cortices. The strongest loadings on sub-factor three were observed in brain activation maps associated with quick motor responses to predominantly auditory stimuli, such as right button and left button responses to auditory cues, and sometimes visual stimuli, such as ‘GO’ stimuli from a stop-signal task or simple visual cues requiring a simple motor response. Interestingly, though activation maps associated with visual stimuli had their strongest loadings on this sub-factor, negative scores were predominantly observed in early visual processing areas. However, the majority of visual activation maps demanded swift motor responses or preparation of motor responses to quickly presented visual cues, which is consistent with positive scores observed in the posterior parietal cortex of the dorsal ‘where’ visual pathway for sub-factor three. Positive scores for sub-factor four outside of the primary auditory cortex were predominantly observed in the inferior frontal gyrus (primarily concentrated in the frontal operculum) and primary visual cortex. The strongest loadings on sub-factor four were observed for activation maps associated with extended processing of complex auditory or visually presented language stimuli, such as sentences, stories, and foreign languages.

Sub-factors two, six, seven and nine were categorized as visual-parietal sub-factors because strong positive scores were observed in the primary visual cortex, inferior and medialtemporal lobe (including the fusiform gyrus), or posterior parietal cortices of these sub-factors. In addition, the activation maps that loaded strongly on each sub-factor involved the presentation of complex visual stimuli, including facial stimuli for sub-factor two, food for sub-factor six, mental manipulation of complex objects and symbols for sub-factor seven, and motor movements in response to facial (same stimuli in sub-factor two) or object stimuli for sub-factor nine. Differences in the pattern of factor scores associated with each sub-factor were also present. For example, positive scores in sub-factor seven extended into the posterior parietal cortex, encompassing both the medial posterior parietal cortex that included the P/PCC region of the DMN, and the superior parietal cortex, consistent with the complex mental and spatial processing associated with the fluid reasoning processes modeled by the paper folding, letter set, and matrix reasoning activation maps.

Sub-factors five, eight and ten were classified as pre-frontal sub-factors given strong positive scores were predominantly observed in lateral and medial prefrontal cortices. The brain activation maps that loaded strongly on these sub-factors involved cognitive processing of visual stimuli, including active viewing of complex visual stimuli for sub-factor five, working-memory and complex cognitive processing processing of visual stimuli (2-back task, mental subtraction, relational comparison, and gambling task) for sub-factor eight, and vocabulary processing of visually presented words for sub-factor ten. Interestingly, the pattern of positive scores in the lateral prefrontal cortices (DLPFC and inferior frontal gyrus) for sub-factors five and ten were strongly lateralized. In particular, positive scores were predominantly in the right lateral prefrontal cortices for sub-factor five and left lateral prefrontal cortices for sub-factor ten, which may reflect hemispheric asymmetries in visuo-spatial and language processing in the right and left hemispheres, respectively (Geschwind, 1970; Shulman et al., 2010). Of note, no positive scores were observed in the medial surface of the primary visual cortex for sub-factor five, though most activation maps loading on this sub-factor were associated with processing of complex visual stimuli. However, positive scores were observed in the lateral occipital and medial temporal areas (regions comprising the ventral ‘where’ pathway), consistent with the complex nature of the visual stimuli.

**Figure 7.**
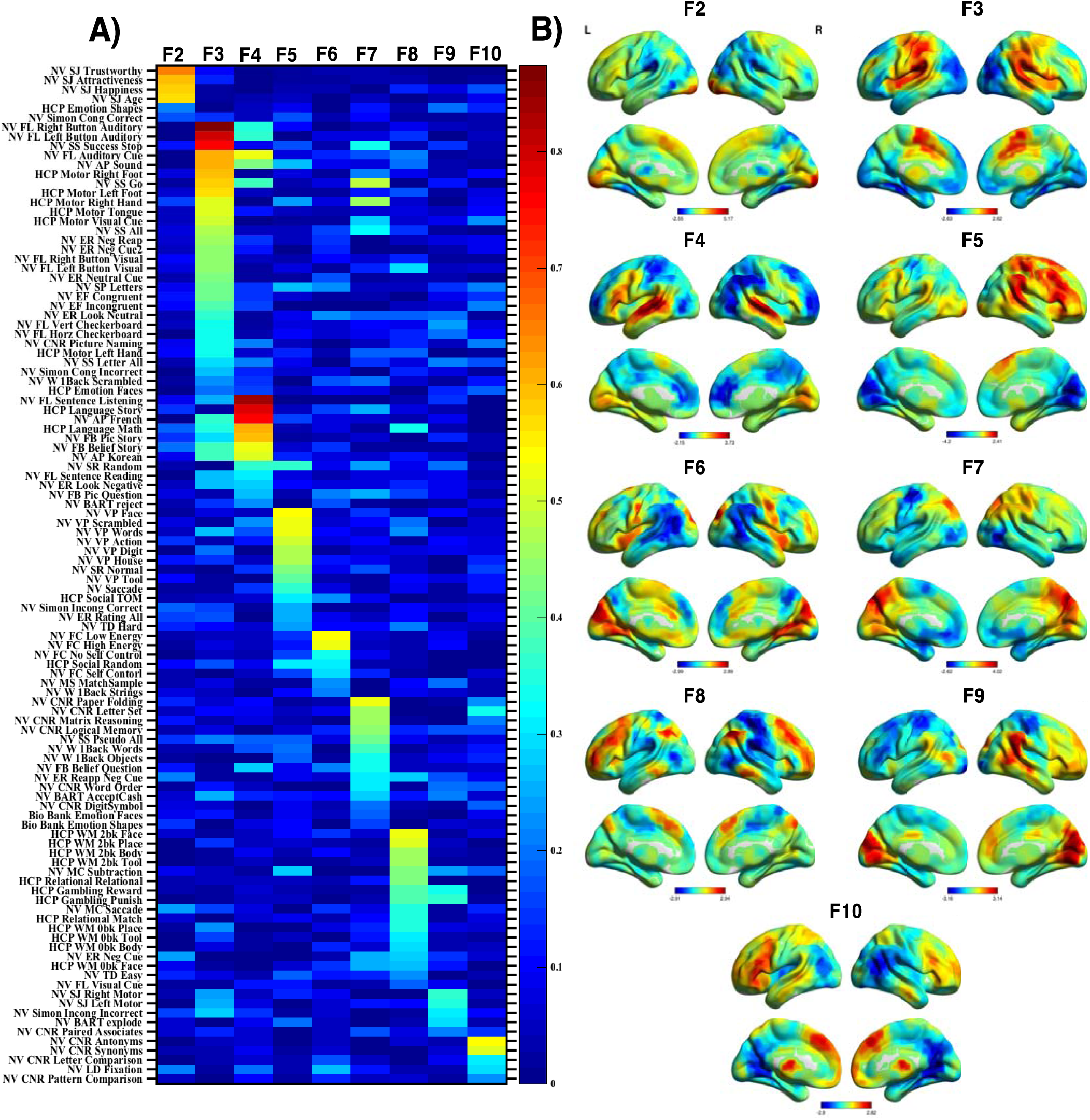
Sub-Factor Loadings and Factor Score Visualization. **A)** The loading of each activation map on each sub-factor, which represents the association between each map and each sub-factor, controlling for the association with other sub-factors. The activation map labels are displayed along the vertical axis and the nine sub-factors (F2-F10) are displayed along the horizontal axis. Factor loadings were represented by a heat map (warmer colors represent stronger loadings and cooler colors represent weaker loadings). For every activation map, the strongest factor loading associated with that map was significant (*p* <.05). **B)** Factor scores associated with each sub-factor were placed in MNI coordinates and visualized on a surface MNI template (warmer colors represent positive scores and cooler colors represent negative factor scores).

### Bi-Factor/Nested Factor Model versus Non-Nested Factor Model

In order to test whether an alternative non-nested factor model fit the data equally well, we used a confirmatory factor analysis (CFA) approach that measured the overall fit of the alternative factor model compared to the bi-factor model. We used a CFA approach that involved comparing the overall fit to the data of a confirmatory bi-factor/nested model derived from the exploratory bi-factor analysis (10 factor model with one general factor and nine sub-factors) and an alternative non-nested factor analysis model constructed from the results of a standard exploratory factor analysis applied to the same set of brain activation maps. The results for all fit indices revealed that the bi-factor model (*RMSEA*=0.16, *CFI*=0.504, *TLI*=0.48, *SRMR=*0.09) had greater overall fit to the data compared to the non-nested standard exploratory factor analysis model (*RMSEA*=0.171, *CFI*=0.415, *TLI*=0.4, *SRMR=*0.158). In addition, to ensure results were not driven by sampling, preprocessing, or scanner-site differences between different collections of brain activation maps, the confirmatory bi-factor/nested model derived from the exploratory bi-factor analysis was also compared with an alternative non-nested study/collection-specific confirmatory model with orthogonal factors representing each study/collection and loadings on each factor from activation maps within that study/collection. The results for all fit indices revealed that the bi-factor model also had greater overall fit to the data than the non-nested study-specific factor model (*RMSEA*=0.175, *CFI*=0.388, *TLI*=0.376, *SRMR=*0.463).

## Discussion

The central goal of this study was to test a hypothesized framework of functional organization of the human brain. We predicted a nested structure of brain activation patterns with a single domain-general brain activation/de-activation pattern, corresponding to the canonical task-positive/task-negative activity pattern, and various sub-groupings of brain-activation patterns that represent different manifestations of the canonical pattern. Specifically, we used the bi-factor analysis model to estimate a nested factor structure involving a single general, overarching factor and various sub-factors within the general factor that underlie the common associations among the brain activation maps. Consistent with the nested factor structure, the single general factor explained the majority of the variance (52.37%) in the collection of brain activation maps, and 94 out of 108 maps had significant loadings on the general factor. In addition to the general factor, were nine sub-factors, associated with various groupings of task demands, that explained another 31.02% of variance in the collection of brain activation maps. Importantly, the nested-factor model discovered provided a better overall fit to the data than an alternative, data-driven non-nested factor model.

The conjunction maps of common activation and de-activation across activation maps reveal that the classical ‘task-positive’ and ‘task-negative’ areas of the brain were the most consistently activated and deactivated areas of the brain across the full set of activation maps, respectively. In line with prior reports demonstrating that the pMPFC is the most consistently activated region across fMRI task studies (Behrens et al., 2013), we found that the pMPFC was activated in over 90% of the activation maps examined. Interestingly, the bilateral fusiform gyri were also found to be as consistently activated as the pMPFC. Another area of high positive scores outside of the traditional ‘task-positive’ areas included the primary motor cortices, possibly corresponding to motor movements from participants in response to experimental stimuli. In addition, de-activation in the classical ‘task-negative’ areas (Raichle et al., 2001), such as the MPFC, precuneus/PCC and bilateral IPC, was on average, much less consistent across the activation maps compared with the ‘task-positive’ areas, indicating that the pattern of de-activation was more variable across different task demands. Of note, none of the brain activation maps exhibited strong activation in the default-mode network. This may be explained by the fact that many reports of default-mode activations during task performance in the literature (Summerfield et al., 2010; Gerlach et al., 2011; Ellamil et al., 2012) are the results of subtraction contrasts, demonstrating differences in activation/de-activation between two modeled conditions (typically an active and control condition), which may result in positive ‘activations’ under circumstances in which the default-mode de-activates less to the active condition compared to the control condition.

While the brain activation maps in the current study exhibited highly consistent patterns of activation/de-activation in certain areas of the cortex, the bi-factor analysis results revealed a more complicated structure of task-positive/task-negative activity patterns not observed in the conjunction analysis. In particular, the bi-factor analysis results are consistent with a bi-factor or nested structure of brain function with a task-general activity pattern that presents in different patterns depending on the task context. Previous literature (Raichle et al., 2001; Fox et al., 2005b; Duncan, 2010; Vanhaudenhuyse et al., 2011; Fedorenko et al., 2013) has focused on a circumscribed collection of lateral and medial frontal and parietal regions as task-positive regions and task-negative regions, respectively. But rather than one canonical task-positive activation and task-negative de-activation, the current results suggest that there are in fact a plurality of manifestations of the traditional task-positive/task-negative pattern depending on the degree of cognitive processing and dominant external stimuli. In fact, frontal and parietal regions most commonly attributed to task-positive and task-negative networks appeared across all sub-factors: positive scores in the pMPFC were observed across all sub-factors, and the DLPFC and AI were observed across most sub-factors; negative scores either in MPFC or P/PCC were observed across all sub-factors.

We suggest that the current results of the bi-factor analysis offer a starting point for a general ontology of psychological categories incorporating data-driven analyses of fMRI data. The canonical task-positive/task-negative pattern of task activity has been hypothesized to indicate a domain-general cognitive process corresponding to ‘focused awareness’ (Hugdahl et al., 2015b) or ‘attentional episodes’ (Duncan, 2010). The domain-general factor encompassed a variety of cognitive processes from several traditionally studied task domains: cognitive control, emotional regulation, selective attention, logical reasoning, short-term memory, mathematical calculation, decision-making, inhibition, language processing, theory of mind and more. The wide range of task domains that significantly load on the general factor strongly suggest a domain-general cognitive process represented by the activation/de-activation patterns, which may indeed be described as a process of ‘focused awareness’, ‘attentional episodes’, or ‘effortful control’ present across all these task states. In addition, the brain activation maps that did not have significant loadings on the general factor involved routine motor responses to simple visual or auditory cues and listening to stories or sentences presented in the auditory modality. It may be that the task demands associated with these activation maps did not require the sort of ‘awareness’ or ‘attention’ associated with the other task demands, but rather involved rote motor responses to simple visual or auditory stimuli or passive listening to sentences. Thus, the cognitive process represented by this domain-general factor may not extend to contexts of automatic, routine response patterns, or passive perception of external stimuli.

Analysis of the sub-factors suggest that what differentiates one form of ‘attention’ or ‘awareness’ from another is the *context* within which it occurs. In other words, rather than one form of ‘awareness’ or ‘attention,’ we suggest multiple types of ‘focused awareness’ and ‘attention episodes,’ depending on the environmental context. For example, a possible interpretation of sub-factors three and four is an *auditory* ‘focused awareness’ or ‘attentional episode’ involving goal-directed responses to auditory stimuli. A possible interpretation of subfactor seven is a visuo-spatial ‘focused awareness’ involving goal-directed attention and mental manipulation of visuospatial stimuli. A possible interpretation of sub-factor eight is a short-term memory ‘focused awareness’ that involves goal-directed attention towards short-term memory traces for subsequent responses. We suggest that the dominant organizing feature of task-activation/de-activation patterns in response to task onsets and offset is not the *content* of the stimulus in and of itself, but the *context* in which that stimulus is presented. We define context in two ways: the nature of the cognitive processing required by the task, such as high-level mental processing versus simple motor responses to visual cues, and the modality of the stimulus presented, such as visual presentation versus auditory presentation. Thus, for sub-factor five, activation maps associated with a variety of types of visual stimuli were presented to participants, but all required attentive processing of visually complex stimuli: determination of whether the category of visual stimulus was presented one-back (Pinel et al., 2015), or determination of whether visual shapes are moving intentionally (theory of mind task). Thus, any task demands that required any sustained attention to, or focused awareness on, experimentally presented stimuli should activate the brain areas common across the sub-factors, but the context in which the stimuli were presented distinguishes this instance of ‘sustained attention’ from another instance.

One particular advantage of using unthresholded statistical images to examine large-scale patterns of brain activity is the additional information regarding task ‘de-activation.’ When researchers examine brain activity in response to task onsets and offsets, these images are normally *positively* thresholded. However, while this data reduction step is informative and crucial for controlling false-positives in single sample studies, much information is lost, including information regarding reduced brain activity to task onsets and offsets, which may be important in understanding the cognitive processes at work in each task condition. In fact, as demonstrated in the factor analytic approach here, negative score patterns can actually distinguish between the different factors that have relatively similar positive score patterns.

## Limitations

While a large variety of brain activation maps from several task domains were examined in this study, the full space of psychological functioning was not covered. An advantage of the analytic approach here involved the use of full-information unthresholded statistic images from a large sample of participants to estimate latent factors, rather than using peak-activation coordinates from online databases. However, the limited availability of unthresholded individual and group statistic images restricts the amount of possible task comparisons. We believe that future studies with larger task comparisons from online resources, such as NeuroVault (Gorgolewski et al., 2015) will be able to achieve a greater sampling of task domains, and thus, a more precise estimation of possible latent factors.

## Conclusion

We propose that a central organizing feature of brain function is a bi-factor or nested factor structure involving a general, over-arching factor and several sub-factors, representing a ‘focused awareness’ and ‘attention allocation’ process that is differentiated by the context in which the process occurs. We take as an assumption, which we believe is shared by most cognitive neuroscientists (but see Bennett and Hacker, 2003; Uttal, 2003), that task fMRI studies are relevant for the delineation of an adequate model of psychological processes (Anderson, 2015). Our approach in this study can be seen as a part of the ongoing project to define a cognitive ontology, or a taxonomy of cognitive processes, in light of neurobiological evidence (Klein, 2012; Anderson, 2015; Poldrack and Yarkoni, 2016). A revised cognitive ontology established on current neurobiological evidence is considered a crucial step in the progress of cognitive neuroscience (Poldrack and Yarkoni, 2016). We also believe that a theory of cognitive processes firmly grounded in neurobiological data would greatly benefit the development of neural and behavioral models of psychiatric and developmental disorders. Such an understanding would greatly support the path towards more reliable clinical diagnoses grounded in neurobiological data (Cuthbert and Insel, 2013).

## Acknowledgments

This work was supported by the National Institute of Mental Health [R01MH107549], a Slifka/Ritvo Innovation in Autism Research Award from the International Society for Autism Research, and a NARSAD Young Investigator Award to LQU.

